# Unique contributions of sensory discrimination and gamma synchronization deficits to cognitive, clinical, and psychosocial functional impairments in schizophrenia

**DOI:** 10.1101/2020.07.19.211193

**Authors:** Daisuke Koshiyama, Makoto Miyakoshi, Michael L. Thomas, Yash B. Joshi, Juan L. Molina, Kumiko Tanaka-Koshiyama, Joyce Sprock, David L. Braff, Neal R. Swerdlow, Gregory A. Light

## Abstract

**Background:** Schizophrenia patients show widespread deficits in neurocognitive, clinical and psychosocial functioning. Mismatch negativity (MMN) and gamma-band auditory steady-state response (ASSR) are robust biomarkers for domains of neuropsychiatric disorders that are impaired in schizophrenia patients and are separately associated with cognitive dysfunction, negative symptom severity and psychosocial disability. Although these measures of early auditory information processing are conceptually linked, it is unclear, whether these measures are redundant or account for unique variance in important outcome measures. In this study, we aimed to determine whether MMN and gamma-band ASSR are associated with cognitive, clinical, and functional variables and, if so, whether they account for shared vs. non-shared variance in those important domains.

**Methods:** Multiple regression analyses with MMN, gamma-band ASSR and clinical measures were performed in large cohorts of schizophrenia outpatients (N=428) and healthy comparison subjects (N=283).

**Results:** Reduced MMN (*d* = 0.67), gamma-band ASSR (*d* = –0.40), and lower cognitive function were confirmed in schizophrenia patients compared to healthy comparison subjects. Regression analyses revealed that both MMN and gamma-band ASSR have significant unique associations with tasks measuring of working memory, and daily functioning in schizophrenia patients.

**Conclusion:** These findings suggest that MMN and ASSR measures are non-redundant and complementary measures. Studies are needed to clarify the neural substrates of MMN and gamma-band ASSR in order to improve our understanding of pathophysiology of schizophrenia and accelerate their use in the development of novel therapeutic interventions.

## 1. Introduction

Patients with schizophrenia show widespread deficits in cognitive, clinical, and psychosocial functioning (Light et al., 2015; Owen et al., 2016; Thomas et al., 2017; van Os and Kapur, 2009). Mismatch negativity (MMN) and the gamma band auditory steady state response (ASSR) are neurophysiological indices related to the pathophysiology of schizophrenia and cognitive dysfunction. Moreover, they are increasingly used as translational biomarkers in the context of the development of novel therapeutics (Braff and Light, 2004; Hochberger et al., 2019; Javitt et al., 2008; Joshi et al., 2018; Kozono et al., 2019; Lavoie et al., 2008; Light and Swerdlow, 2015; Light et al., 2015; Perez et al., 2017; Swerdlow et al., 2016).

MMN is an event-related potential (ERP) measure of sensory discrimination processing. MMN is typically measured in the context of a passive auditory oddball paradigm where a series of identical standard tones (e.g. 90%) is interspersed with less frequent (e.g. 10%) “oddball” stimuli that differ in some physical characteristic such as stimulus duration, pitch or loudness. The MMN is a negative-going peak reflecting the differences between scalp-level ERP responses to deviant vs. standard stimuli measured across a 135–205-ms time window and thought to reflect an automatic deviance detection process. Several studies have reported large effect sizes for MMN differences between schizophrenia patients and controls using electroencephalography (EEG; Hermens et al., 2010; Javitt et al., 1995; Koshiyama et al., 2017, 2018b, 2018c; Light and Braff, 2005; Light et al., 2015; Rasser et al., 2011; Salisbury et al., 2017; Wynn et al., 2010) or magnetoencephalography (Braeutigam et al., 2018; Yamasue et al., 2004), and these findings have been confirmed by several meta-analyses (Erickson et al., 2016; Umbricht and Krljes, 2005); These impairments are among the most consistently replicated findings in schizophrenia research (Keshavan et al., 2008; Näätänen et al., 2019). Many studies have also identified important associations between MMN and cognition (Koshiyama et al., 2018c; Rissling et al., 2014; Toyomaki et al., 2008; Wynn et al., 2010), negative symptoms (Light et al., 2015) and functional outcomes (Koshiyama et al., 2018c; Light and Braff, 2005; Rissling et al., 2014) in schizophrenia patients.

The gamma-band ASSR is also a robust biomarker that is increasingly studied in neuropsychiatric disorders (Tada et al., 2019a; Uhlhaas and Singer, 2010). Previous studies have found selectively reduced power and synchronization in response to 40-Hz auditory stimulation in schizophrenia patients using EEG (Brenner et al., 2003; Hamm et al., 2015; Hirano et al., 2015; Kirihara et al., 2012; Koshiyama et al., 2018a, 2018b, 2019; Kwon et al., 1999; Light et al., 2006; Spencer et al., 2008; Tada et al., 2016) and MEG (Edgar et al., 2014; Teale et al., 2008; Tsuchimoto et al., 2011; Vierling-Claassen et al., 2008; Wilson et al., 2008). A recent meta-analysis confirmed that the 40-Hz ASSR is a robust index of gamma synchronization deficits in schizophrenia patients (Thuné et al., 2016). While several studies have linked ASSR dysfunction to both cognitive (Light et al., 2006; Tada et al., 2016) and psychosocial functioning (Koshiyama et al., 2018a) in schizophrenia, the associations among gamma-band ASSR, neurocognition, clinical symptoms and psychosocial functioning are not fully understood.

MMN and ASSR are both EEG measures of early auditory processing with associations among cognitive, clinical, and functional domains. Indeed, our previous study demonstrated a correlation between reduced MMN amplitude and reduced gamma-band ASSR in schizophrenia patients (Koshiyama et al., 2018b). However, it is not yet clear to what degree they may be shared or may account for unique variance in important outcome measures. In this study, we aimed to determine whether MMN and gamma-band ASSR are associated with neurocognitive, clinical, and functional domains and, if so, whether they account for specific variance in these important domains.

## 2. Material and methods

### 2.1. Subjects

Participants included healthy comparison subjects (n=283) and patients diagnosed with schizophrenia (n=428). The subsets of these patients were reported in previous studies. Primary ASSR deficits were originally reported in a subset of participants (n=188, healthy comparison subjects; n=234, schizophrenia patients; Kirihara et al., 2012). MMN deficits as subsets were published in a couple different contexts (n=247, healthy comparison subjects; n=410, schizophrenia patients; Takahashi et al., 2013). Patients were diagnosed based on a clinical interview using a modified version of the Structured Clinical Interview for DSM-IV-TR. Patients were recruited from community residential facilities and via clinician referral. Of the 428 patients, 388 were stabilized on antipsychotic medications (three patients had no medication data). Healthy comparison subjects were recruited through internet advertisements. Exclusion criteria included an inability to understand the consent processes and/or provide consent or assent, not being a fluent English speaker, previous significant head injury with loss of consciousness, neurological illness, severe systemic illness, or current mania. Written informed consent was obtained from each subject. The Institutional Review Board of University of California San Diego approved all experimental procedures (071128, 071831 and 170147).

### 2.2. Cognition, clinical symptoms and functional outcome assessment

Working memory was evaluated using Letter-Number Sequencing (LNS) with higher scores indicating greater ability (Crowe, 2000; standard scores were employed). Verbal learning performance was measured using California verbal learning test (CVLT) second edition using total correct scores from the Total Learning (list A trials 1–5) with higher scores indicating greater ability (Delis et al., 2000; Stone et al., 2011; standard scores were employed). Executive function was assessed using the number of perseverative responses obtained in the Wisconsin Card Sorting Test (WCST) with lower scores indicating greater ability (Berg, 1948; Grant and Berg, 1948). Clinical symptoms were assessed with the Scale for the Assessment of Negative Symptoms (SANS; Andreasen, 1984). Functional outcomes were measured using the Scale of Functioning (SOF; Rapaport et al., 1996) and the Global Assessment of Functioning scale (GAF). Higher scores indicate better functioning for the SOF. The GAF is a scale that evaluates the overall level of social adaptation from 0 to 100 scores. A higher score means a higher global function.

### 2.3. Mismatch negativity

Subjects were presented with binaural tones (1-kHz, 85-dB, with 1-ms rise/fall, stimulus onset asynchrony 500 ms) via insert earphones (Aearo Company Auditory Systems, Indianapolis, IN; Model 3A). A duration-deviant auditory oddball paradigm where the deviant stimuli differed in duration was employed following our established procedures (Rissling et al., 2014). Standard (*p* = 0.90, 50-ms duration) and Deviant (*p* = 0.10, 100-ms duration) tones were presented in pseudorandom order with a minimum of 6 Standard stimuli presented between each Deviant stimulus. The MMN amplitude at the Fz was measured using the mean voltage from 135 to 205 ms post stimuli in accordance with previous studies (Koshiyama et al., 2017, 2018b, 2018c, in press). During the MMN and ASSR sessions, participants watched a silent cartoon video.

### 2.4. Gamma band auditory steady-state response

Auditory steady-state stimuli were 1-millisecond, 93-dB clicks presented at 40 Hz in 500-millisecond trains. A block typically contained 200 trains of the clicks with 500-millisecond intervals. The ASSR at the Fz was used for analysis. We performed time-frequency analyses with a short-term Fourier transformation (STFT) and then calculated event-related spectral perturbation (ERSP) as an index of ASSR. We calculated the mean ERSP by averaging the data over stimulation time (0–500 ms) and stimulation frequency (40 Hz: 35–45 Hz) in accordance with previous studies (Koshiyama et al., 2018a, 2018b, 2019).

### 2.5. EEG recording and preprocessing

EEG data were continuously digitized at a rate of 500 Hz (nose reference, forehead ground) using a 40-channel Neuroscan system (Neuroscan Laboratories, El Paso, Texas). The electrode montage was based on standard positions in the International 10–5 electrode system (Oostenveld and Praamstra, 2001) fit to the MNI template head used in EEGLAB, including AFp10 and AFp9 as horizontal EOG channels, IEOG and SEOG above and below the left eye as vertical EOG channels, Fp1, Fp2, F7, F8, Fz, F3, F4, FC1, FC2, FC5, FC6, C3, Cz, C4, CP1, CP2, CP5, CP6, P7, P3, Pz, P4, P8, T7, T8, TP9, TP10, FT9, FT10, PO9, PO10, O1, O2, and Iz. Electrode-to-skin impedance mediated by conductive gel was brought below 4 kΩ. The system acquisition band pass was 0.5–100 Hz. Offline, EEG data were imported to EEGLAB 14.1.2 (Delorme and Makeig, 2004) running under Matlab 2017b (The MathWorks, Natick, MA). Data were high-pass filtered (FIR, Hamming window, cutoff frequency 0.5 Hz, transition bandwidth 0.5).

For data cleaning, EEGLAB plugin *clean_rawdata()* was applied including Artifact Subspace Reconstruction was applied to reduce high-amplitude artifacts (Blum et al., 2019; Chang et al., 2018, 2020; Gabard-Durnam et al., 2018; Kothe and Makeig, 2013; Mullen et al., 2015; Plechawska-Wojcik M, 2019). The parameters used were: flat line removal, 10 s; electrode correlation, 0.7; ASR, 20; window rejection, 0.5. As a result, for the 711 MMN datasets, mean channel rejection rate was 4.2 % (SD 2.4, range 0-15.8). Mean data rejection rate was 3.1% (SD 3.9, range 0-28.7). For the 711 ASSR datasets, mean channel rejection rate was 4.2 % (SD 2.3, range 0-15.8). Mean data rejection rate was 2.1% (SD 3.8, range 0-38.7). The rejected channels were interpolated using EEGLAB’s spline interpolation function. Data were re-referenced to average.

Adaptive mixture ICA (Bell and Sejnowski, 1995; Delorme et al., 2012; Palmer J, 2016; Palmer J, 2008; Palmer and Rosa, 2006) was applied to the preprocessed scalp recording data to obtain temporally maximally independent components. For scalp topography of each independent component derived, equivalent current dipole was estimated using Fieldtrip functions (Oostenveld et al., 2011). For scalp topographies more suitable for symmetrical bilateral dipoles, two symmetrical dipoles were estimated (Piazza C, 2016).

To select non-artifact ICs among all types of ICs, EEGLAB plugin *ICLabel()* was used (Pion-Tonachini et al., 2019). Those ICs with any of the following label that showed label probability >= 0.7 was rejected: ‘muscle’, ‘eye’, ‘heart’, ‘line noise’, and ‘channel noise’. The purpose of this approach was to approximate the process to the conventional manual artifact rejection procedure in which ICA is used to identify artifacts for rejection. Compared with the alternative approach using label probability of ‘brain’ as a single criterion, the current approach is more tolerant to marginal and mixed evidence of brain-origin signals hence may be more sensitive as such.

As a result, for MMN, the mean number of independent components included for scalp-electrode backprojection was 26.1 (SD 3.9, range 12-36). For ASSR, the mean number of independent components included for backprojection to scalp electrodes was 26.8 (SD 3.9, range 13-35). This is equivalent to the rejection of 12-13 ICs per subject that represent various types of artifacts as described above. In the present study, time-series data associated with the scalp electrode Fz was re-constructed by linear combination of the average 25-27 artifact-free ICs back-projecting from their cortically-resolved source locations. The removal of artifact-laden ICs enhances the robustness of neural source contributions at the level of scalp-level topographies relative to conventional single-trial artifact removal methods (Onton and Makeig, 2006).

### 2.6. Statistical analysis

Independent samples *t-*tests were used to compare schizophrenia patient and healthy comparison subject groups on cognitive function, MMN and gamma-band ASSR. The significance level was set to *p* < 0.05. Cohen’s *d* was used to quantify effect size. In order to examine the contributions of MMN and gamma-band ASSR to predicting cognitive function, negative symptoms and functional outcomes, regression analyses were performed in schizophrenia patients using the following formula:

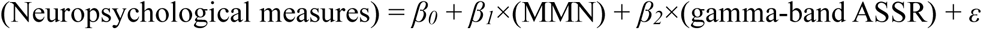

The significance level was set to *p* < 8.3×10^−3^ (0.05/6; 6 neuropsychological measures) adjusted for Bonferroni correction.

## 3. Results

### 3.1. Difference of cognitive function, MMN and gamma-band ASSR between healthy comparison subjects and schizophrenia patients

As shown in **Table 1**, schizophrenia patients showed significant impairments across all cognitive domains (LNS, *d* = –1.17; CVLT, *d* = –1.34; WCST, *d* = 0.77), MMN (*d* = 0.67) and gamma-band ASSR (*d* = –0.40). The average MMN waveforms and the time-course and grand average time-frequency maps for the ERSP at Fz for each group are shown in **Figure 1**.

**Table 1.** Differences of cognition, MMN and gamma-band ASSR between healthy comparison subjects and schizophrenia patients and clinical symptoms and functional outcome in schizophrenia patients.

|  | Healthy comparison subjects |  | Schizophrenia patients |  | Statistics |  |  |  |
| --- | --- | --- | --- | --- | --- | --- | --- | --- |
|  | Mean | SD | Mean | SD | *d* | *t* | *df* | *p* |
| LNS<sup>a</sup> | 10.6 | 2.4 | 7.6 | 2.7 | −1.17 | 15.2 | 705 | 2.4×10<sup>−45*</sup> |
| CVLT<sup>b</sup> | 52.4 | 11.3 | 36.6 | 12.1 | −1.34 | 17.4 | 703 | 7.3×10<sup>−57*</sup> |
| WCST<sup>c</sup> | 11.4 | 8.9 | 22.8 | 17.8 | 0.77 | −10.0 | 701 | 4.3×10<sup>−22*</sup> |
| SANS total<sup>d</sup> |  |  | 14.4 | 4.3 |  |  |  |  |
| SOF total<sup>e</sup> |  |  | 44.3 | 5.3 |  |  |  |  |
| GAF<sup>f</sup> |  |  | 40.9 | 6.5 |  |  |  |  |
| MMN | −1.95 | 1.31 | −1.19 | 1.01 | 0.67 | −8.8 | 709 | 1.3×10<sup>−17*</sup> |
| Gamma-band ASSR | 0.78 | 0.64 | 0.53 | 0.62 | −0.40 | 5.2 | 709 | 3.0×10<sup>−7*</sup> |

Legend: Asterisks indicate statistical significance set at $p < 0.05$ ; *d* indicates effect size of Cohen's *d*; Event-related spectral perturbation is used as an index of gamma-band ASSR; <sup>a</sup>Four subjects have no data of LNS; <sup>b</sup>Six subjects have no data of CVLT; <sup>c</sup>Eight subjects have no data of WCST; <sup>d</sup>Four subjects have no data of SANS; <sup>e</sup>Thirteen subjects have no data of SOF; <sup>f</sup>One subject has no data of GAF.
Abbreviation: SD, standard deviation; LNS, letter-number sequencing; CVLT, California verbal learning test; WCST, Wisconsin card sorting test; SANS, scale for the assessment of negative symptoms; SOF, scale of functioning; GAF, global functioning; MMN, mismatch negativity; ASSR, auditory steady-state response.

**Figure 1.**
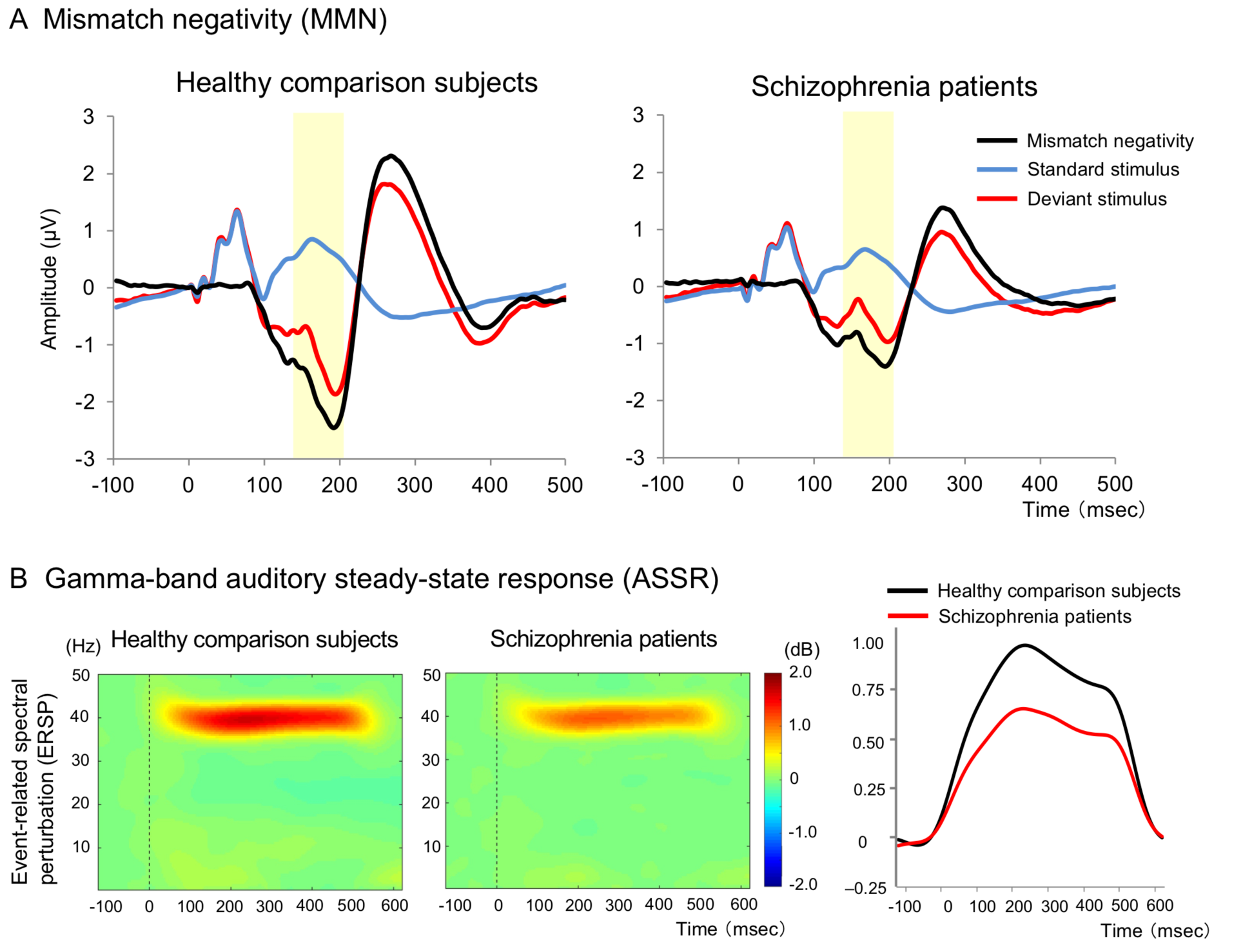
Mismatch negativity (A) and gamma-band auditory steady-state response (B) at Fz. Legend: Yellow shadow indicates mismatch negativity time window (135–205 ms).

### 3.2. Effects of MMN and gamma-band ASSR on cognition, negative symptoms and functional outcomes in schizophrenia patients

MMN and gamma-band ASSR both significantly predicted LNS scores (MMN, *β* = – 0.17; gamma-band ASSR, *β* = 0.15) and SOF total scores (MMN, *β* = –0.23; gamma-band ASSR, *β* = 0.16) in schizophrenia patients (**Table 2**). Additionally, MMN uniquely predicted CVLT scores (*β* = –0.15), WCST scores (*β* = 0.20), SANS total scores (*β* = 0.18) and GAF scores (*β* = –0.17). There were no other significant effects of MMN or gamma-band ASSR on cognition, negative symptoms or functional outcomes.

**Table 2.** Effect of MMN and gamma-band ASSR on cognition, clinical symptoms and functional outcomes in schizophrenia patients.

|  | MMN |  | Gamma-band ASSR |  |
| --- | --- | --- | --- | --- |
| | $\beta$ | $p$ | $\beta$ | $p$ |
| LNS | -0.17 | $5.5 \times 10^{-4*}$ | 0.15 | $2.1 \times 10^{-3*}$ |
| CVLT | -0.15 | $1.6 \times 10^{-3*}$ | 0.12 | $1.8 \times 10^{-2}$ |
| WCST | 0.20 | $4.1 \times 10^{-5*}$ | -0.03 | 0.54 |
| SANS total | 0.18 | $2.8 \times 10^{-4*}$ | -0.12 | $1.8 \times 10^{-2}$ |
| SOF total | -0.23 | $2.5 \times 10^{-6*}$ | 0.16 | $9.1 \times 10^{-4*}$ |
| GAF | -0.17 | $4.6 \times 10^{-4*}$ | 0.05 | 0.27 |

**Legend:** Asterisks indicate statistical significance set at $p < 8.3 \times 10^{-3}$ (0.05/6; 6 neuropsychological measures) adjusted for the Bonferroni correction. $\beta$ is standardized coefficient beta. Event-related spectral perturbation is used as an index of gamma-band ASSR.
Abbreviation: MMN, mismatch negativity; ASSR, auditory steady-state response; LNS, letter-number sequencing; CVLT, California verbal learning test; WCST, Wisconsin card sorting test; SANS, scale for the assessment of negative symptoms; GAF, global functioning.

## 4. Discussion

These results demonstrate that MMN and gamma band ASSR are both reduced in schizophrenia patients and are both associated with cognitive and psychosocial functioning deficits in schizophrenia patients. Importantly, MMN and gamma-band ASSR are non-redundant measures of early auditory information processing and account for unique variance in multiple domains of cognition. MMN was uniquely associated with verbal learning, executive function, and negative symptoms in this cohort of schizophrenia patients.

### 4.1. Relationship to previous MMN studies

The relationships described in this report are consistent with previous separate findings of either MMN or gamma-band ASSR associations with cognition, clinical symptoms and psychosocial impairments. For example, MMN has been associated with working memory (Koshiyama et al., 2018c; Thomas et al., 2017), executive function (Kiang et al., 2007; Toyomaki et al., 2008) and psychosocial impairments (Kawakubo and Kasai, 2006; Koshiyama et al., 2018c; Light and Braff, 2005; Thomas et al., 2017; Wynn et al., 2010) in schizophrenia patients. Consistent with those results, the current findings confirm that MMN is related to a broad range of impairments in neurocognitive and psychosocial domains in schizophrenia. While a previous meta-analysis failed to detect associations between negative symptoms and MMN in schizophrenia patients (Erickson et al., 2017), our previous study investigating MMN in a non-overlapping sample of schizophrenia patients from the Consortium on the Genetics of Schizophrenia (COGS)-2 data (N = 966) identified a significant but modest correlation between MMN amplitude and negative symptoms (Light et al., 2015).

### 4.2. Relationship to previous gamma-band ASSR studies

The significant association between gamma-band ASSR and working memory is consistent with prior findings (Light et al., 2006). We have also previously (Koshiyama et al., 2018a) reported that evoked gamma oscillations predict global symptomatic outcomes after 1-2 years in recent-onset schizophrenia patients who did not show significant ASSR relationships with symptoms at the start of the study. Notably, gamma-band ASSR was not significantly associated with GAF scores in the present study, but significant associations were detected between ASSR and SOF. We speculate that since the SOF exclusively focuses on domains of independent living, social, and instrumental functioning, and that GAF reflects a combination of both clinical symptom severity and psychosocial outcomes (including outcomes more distinctly operationalized in the SOF), ASSR is more sensitive to symptom-insensitive functional outcomes.

### 4.3. Unique contributions of MMN and gamma-band ASSR to cognition and psychosocial functioning

Although both MMN and gamma-band ASSR are neurophysiological indices of early auditory information processing, the present study found that these measures show unique contributions to working memory performance and psychosocial functioning and are thus complementary measures in schizophrenia research. Despite their conceptual similarity as measures of early auditory information processing, the fact that MMN and ASSR are dissociable in their relationship to distinct clinical features suggests that they may reflect different aspects of underlying neurobiology (Tada et al., 2019, in press). These findings suggest that MMN and gamma-band ASSR are complementary biomarkers; combining these biomarkers may help identify clusters of schizophrenia patients with similar but distinct clinical and/or neurophysiologic profiles and may inform future biomarker-guided approaches for therapeutic development.

### 4.4. Limitations

This study should be considered in the context of several limitations. First, this is a cross-sectional cohort study of a heterogeneous sample of schizophrenia patients, the majority of whom were receiving complex medication regimens. As is the case for most large-scale studies of schizophrenia patients, the medication, psychosocial environments, and other important factors that could potentially influence brain function were not experimentally controlled. Second, the relatively modest effect sizes in this study may be due to the substantial heterogeneity of the sample. Third, scalp level EEG data at Fz was used in this study, which has known source contributions from multiple brain regions (Rissling et al., 2014). It is possible that future studies investigating source level analyses representing the independent contributions from individual brain regions may better clarify the relationship between MMN and ASSR, and their cognitive, clinical and functional correlates. Last, schizophrenia patients in this study had a “chronic” illness; results therefore may not generalize to at risk or early-illness psychosis patients.

### 4.5. Conclusion

In this study, we showed significant correlations of both reduced MMN amplitude and reduced gamma-band ASSR with working memory deficits and daily functioning impairment in schizophrenia patients. These results demonstrated that MMN and gamma-band ASSR account for unique and additive portions of variance in important outcome variables and are thus complementary rather than redundant associated measures of early auditory information processing. Furthermore, reduced MMN amplitude has a significant association with lower verbal learning, executive function and negative symptoms in schizophrenia patients. Investigation of the neural substrates of MMN and gamma-band ASSR and their associations with cognitive dysfunction, clinical symptoms and functional impairment will strengthen the utility of MMN and gamma-band ASSR as important biomarkers for clarifying the pathophysiology of schizophrenia and the development of novel therapeutic interventions.

## Acknowledgements

We gratefully acknowledge all the participants of this study.

## Author contributions

J. Sprock, D. Braff, N. Swerdlow and G. Light collected the data. D. Koshiyama and M. Miyakoshi analyzed the data. D. Koshiyama, M. Miyakoshi, M. Thomas, Y. Joshi, J. Molina, K. Tanaka-Koshiyama, D. Braff, N. Swerdlow and G. Light interpreted the results. D. Koshiyama and G. Light designed the study. G. Light supervised all aspects of collection, analysis, and interpretation of the data. D. Koshiyama, M. Miyakoshi and G. Light wrote original manuscript. M. Thomas, Y. Joshi, J. Molina, K. Tanaka-Koshiyama, J. Sprock, D. Braff and N. Swerdlow reviewed and edited the manuscript. All authors contributed to and approved the final manuscript.

## References

Andreasen, N.C., 1984. The scale for the assessment of negative symptoms (SANS). University of Iowa, Iowa CIty.

Bell, A.J., Sejnowski, T.J., 1995. An information-maximization approach to blind separation and blind deconvolution. Neural Comput 7(6), 1129–1159.

Berg, E.A., 1948. A simple objective technique for measuring flexibility in thinking. J Gen Psychol 39, 15–22.

Blum, S., Jacobsen, N.S.J., Bleichner, M.G., Debener, S., 2019. A Riemannian Modification of Artifact Subspace Reconstruction for EEG Artifact Handling. Front Hum Neurosci 13, 141.

Braeutigam, S., Dima, D., Frangou, S., James, A., 2018. Dissociable auditory mismatch response and connectivity patterns in adolescents with schizophrenia and adolescents with bipolar disorder with psychosis: A magnetoencephalography study. Schizophr Res 193, 313–318.

Braff, D.L., Light, G.A., 2004. Preattentional and attentional cognitive deficits as targets for treating schizophrenia. Psychopharmacology (Berl) 174(1), 75–85.

Brenner, C.A., Sporns, O., Lysaker, P.H., O’Donnell, B.F., 2003. EEG synchronization to modulated auditory tones in schizophrenia, schizoaffective disorder, and schizotypal personality disorder. Am J Psychiatry 160(12), 2238–2240.

Chang, C.Y., Hsu, S.H., Pion-Tonachini, L., Jung, T.P., 2018. Evaluation of Artifact Subspace Reconstruction for Automatic EEG Artifact Removal. Conf Proc IEEE Eng Med Biol Soc 2018, 1242–1245.

Chang, C.Y., Hsu, S.H., Pion-Tonachini, L., Jung, T.P., 2020. Evaluation of Artifact Subspace Reconstruction for Automatic Artifact Components Removal in Multi-channel EEG Recordings. IEEE Trans Biomed Eng 67, 1114–1121.

Crowe, S.F., 2000. Does the letter number sequencing task measure anything more than digit span? Assessment 7(2), 113–117.

Delis, D.C., Kramer, J.H., Kaplan, E., Ober, B.A., 2000. California Verbal Learning Test, second edition. Psychological Corporation, San Antonio, TX.

Delorme, A., Makeig, S., 2004. EEGLAB: an open source toolbox for analysis of single-trial EEG dynamics including independent component analysis. J Neurosci Methods 134(1), 9–21.

Delorme, A., Palmer, J., Onton, J., Oostenveld, R., Makeig, S., 2012. Independent EEG sources are dipolar. PLoS One 7(2), e30135.

Edgar, J.C., Chen, Y.H., Lanza, M., Howell, B., Chow, V.Y., Heiken, K., Liu, S., Wootton, C., Hunter, M.A., Huang, M., Miller, G.A., Canive, J.M., 2014. Cortical thickness as a contributor to abnormal oscillations in schizophrenia? Neuroimage Clin 4, 122–129.

Erickson, M.A., Albrecht, M., Ruffle, A., Fleming, L., Corlett, P., Gold, J., 2017. No association between symptom severity and MMN impairment in schizophrenia: A meta-analytic approach. Schizophr Res Cogn 9, 13–17.

Erickson, M.A., Ruffle, A., Gold, J.M., 2016. A meta-analysis of mismatch negativity in schizophrenia: from clinical risk to disease specificity and progression. Biol Psychiatry 79(12), 980–987.

Gabard-Durnam, L.J., Mendez Leal, A.S., Wilkinson, C.L., Levin, A.R., 2018. The Harvard Automated Processing Pipeline for Electroencephalography (HAPPE): Standardized Processing Software for Developmental and High-Artifact Data. Front Neurosci 12, 97.

Grant, D.A., Berg, E.A., 1948. A behavioral analysis of degree of reinforcement and ease of shifting to new responses in a Weigl-type card-sorting problem. J Exp Psychol 38(4), 404–411.

Hamm, J.P., Bobilev, A.M., Hayrynen, L.K., Hudgens-Haney, M.E., Oliver, W.T., Parker, D.A., McDowell, J.E., Buckley, P.A., Clementz, B.A., 2015. Stimulus train duration but not attention moderates gamma-band entrainment abnormalities in schizophrenia. Schizophr Res 165(1), 97–102.

Hermens, D.F., Ward, P.B., Hodge, M.A., Kaur, M., Naismith, S.L., Hickie, I.B., 2010. Impaired MMN/P3a complex in first-episode psychosis: cognitive and psychosocial associations. Prog Neuropsychopharmacol Biol Psychiatry 34(6), 822–829.

Hirano, Y., Oribe, N., Kanba, S., Onitsuka, T., Nestor, P.G., Spencer, K.M., 2015. Spontaneous Gamma Activity in Schizophrenia. JAMA Psychiatry 72(8), 813–821.

Hochberger, W.C., Joshi, Y.B., Thomas, M.L., Zhang, W., Bismark, A.W., Treichler, E.B.H., Tarasenko, M., Nungaray, J., Sprock, J., Cardoso, L., Swerdlow, N., Light, G.A., 2019. Neurophysiologic measures of target engagement predict response to auditory-based cognitive training in treatment refractory schizophrenia. Neuropsychopharmacology 44(3), 606–612.

Javitt, D.C., Doneshka, P., Grochowski, S., Ritter, W., 1995. Impaired mismatch negativity generation reflects widespread dysfunction of working memory in schizophrenia. Arch Gen Psychiatry 52(7), 550–558.

Javitt, D.C., Spencer, K.M., Thaker, G.K., Winterer, G., Hajos, M., 2008. Neurophysiological biomarkers for drug development in schizophrenia. Nat Rev Drug Discov 7(1), 68–83.

Joshi, Y.B., Breitenstein, B., Tarasenko, M., Thomas, M.L., Chang, W.L., Sprock, J., Sharp, R.F., Light, G.A., 2018. Mismatch negativity impairment is associated with deficits in identifying real-world environmental sounds in schizophrenia. Schizophr Res 191, 5–9.

Kawakubo, Y., Kasai, K., 2006. Support for an association between mismatch negativity and social functioning in schizophrenia. Prog Neuropsychopharmacol Biol Psychiatry 30(7), 1367–1368.

Keshavan, M.S., Tandon, R., Boutros, N.N., Nasrallah, H.A., 2008. Schizophrenia, “just the facts”: what we know in 2008 Part 3: neurobiology. Schizophr Res 106(2-3), 89–107.

Kiang, M., Light, G.A., Prugh, J., Coulson, S., Braff, D.L., Kutas, M., 2007. Cognitive, neurophysiological, and functional correlates of proverb interpretation abnormalities in schizophrenia. J Int Neuropsychol Soc 13(4), 653–663.

Kirihara, K., Rissling, A.J., Swerdlow, N.R., Braff, D.L., Light, G.A., 2012. Hierarchical organization of gamma and theta oscillatory dynamics in schizophrenia. Biol Psychiatry 71(10), 873–880.

Koshiyama, D., Kirihara, K., Tada, M., Nagai, T., Fujiok, M., Usui, K., Araki, T., Kasai, K., in press. Reduced auditory mismatch negativity reflects impaired deviance detection in schizophrenia. Schizophr Bull.

Koshiyama, D., Kirihara, K., Tada, M., Nagai, T., Fujioka, M., Ichikawa, E., Ohta, K., Tani, M., Tsuchiya, M., Kanehara, A., Morita, K., Sawada, K., Matsuoka, J., Satomura, Y., Koike, S., Suga, M., Araki, T., Kasai, K., 2018a. Auditory gamma oscillations predict global symptomatic outcome in the early stages of psychosis: A longitudinal investigation. Clin Neurophysiol 129(11), 2268–2275.

Koshiyama, D., Kirihara, K., Tada, M., Nagai, T., Fujioka, M., Ichikawa, E., Ohta, K., Tani, M., Tsuchiya, M., Kanehara, A., Morita, K., Sawada, K., Matsuoka, J., Satomura, Y., Koike, S., Suga, M., Araki, T., Kasai, K., 2018b. Electrophysiological evidence for abnormal glutamate-GABA association following psychosis onset. Transl Psychiatry 8(1), 211.

Koshiyama, D., Kirihara, K., Tada, M., Nagai, T., Fujioka, M., Koike, S., Suga, M., Araki, T., Kasai, K., 2018c. Association between mismatch negativity and global functioning is specific to duration deviance in early stages of psychosis. Schizophr Res 195, 378–384.

Koshiyama, D., Kirihara, K., Tada, M., Nagai, T., Fujioka, M., Usui, K., Koike, S., Suga, M., Araki, T., Hashimoto, K., Kasai, K., 2019. Gamma-band auditory steady-state response is associated with plasma levels of d-serine in schizophrenia: An exploratory study. Schizophr Res 208, 467–469.

Koshiyama, D., Kirihara, K., Tada, M., Nagai, T., Koike, S., Suga, M., Araki, T., Kasai, K., 2017. Duration and frequency mismatch negativity shows no progressive reduction in early stages of psychosis. Schizophr Res 190, 32–38.

Kothe, C.A., Makeig, S., 2013. BCILAB: a platform for brain-computer interface development. J Neural Eng 10(5), 056014.

Kozono, N., Honda, S., Tada, M., Kirihara, K., Zhao, Z., Jinde, S., Uka, T., Yamada, H., Matsumoto, M., Kasai, K., Mihara, T., 2019. Auditory Steady State Response; nature and utility as a translational science tool. Sci Rep 9(1), 8454.

Kwon, J.S., O’Donnell, B.F., Wallenstein, G.V., Greene, R.W., Hirayasu, Y., Nestor, P.G., Hasselmo, M.E., Potts, G.F., Shenton, M.E., McCarley, R.W., 1999. Gamma frequency-range abnormalities to auditory stimulation in schizophrenia. Arch Gen Psychiatry 56(11), 1001–1005.

Lavoie, S., Murray, M.M., Deppen, P., Knyazeva, M.G., Berk, M., Boulat, O., Bovet, P., Bush, A.I., Conus, P., Copolov, D., Fornari, E., Meuli, R., Solida, A., Vianin, P., Cuenod, M., Buclin, T., Do, K.Q., 2008. Glutathione precursor, N-acetyl-cysteine, improves mismatch negativity in schizophrenia patients. Neuropsychopharmacology 33(9), 2187–2199.

Light, G.A., Braff, D.L., 2005. Mismatch negativity deficits are associated with poor functioning in schizophrenia patients. Arch Gen Psychiatry 62(2), 127–136.

Light, G.A., Hsu, J.L., Hsieh, M.H., Meyer-Gomes, K., Sprock, J., Swerdlow, N.R., Braff, D.L., 2006. Gamma band oscillations reveal neural network cortical coherence dysfunction in schizophrenia patients. Biol Psychiatry 60(11), 1231–1240.

Light, G.A., Swerdlow, N.R., 2015. Future clinical uses of neurophysiological biomarkers to predict and monitor treatment response for schizophrenia. Ann N Y Acad Sci 1344, 105–119.

Light, G.A., Swerdlow, N.R., Thomas, M.L., Calkins, M.E., Green, M.F., Greenwood, T.A., Gur, R.E., Gur, R.C., Lazzeroni, L.C., Nuechterlein, K.H., Pela, M., Radant, A.D., Seidman, L.J., Sharp, R.F., Siever, L.J., Silverman, J.M., Sprock, J., Stone, W.S., Sugar, C.A., Tsuang, D.W., Tsuang, M.T., Braff, D.L., Turetsky, B.I., 2015. Validation of mismatch negativity and P3a for use in multi-site studies of schizophrenia: characterization of demographic, clinical, cognitive, and functional correlates in COGS-2. Schizophr Res 163(1-3), 63–72.

Mullen, T.R., Kothe, C.A., Chi, Y.M., Ojeda, A., Kerth, T., Makeig, S., Jung, T.P., Cauwenberghs, G., 2015. Real-Time Neuroimaging and Cognitive Monitoring Using Wearable Dry EEG. IEEE Trans Biomed Eng 62(11), 2553–2567.

Näätänen, R., Kujala, T., Light, G., 2019. The Mismatch Negativity: A Window to the Brain. Oxford University Press, Oxford.

Onton, J., Makeig, S., 2006. Information-based modeling of event-related brain dynamics. Prog Brain Res 159, 99–120.

Oostenveld, R., Fries, P., Maris, E., Schoffelen, J.M., 2011. FieldTrip: Open source software for advanced analysis of MEG, EEG, and invasive electrophysiological data. Comput Intell Neurosci 2011, 156869.

Oostenveld, R., Praamstra, P., 2001. The five percent electrode system for high-resolution EEG and ERP measurements. Clin Neurophysiol 112(4), 713–719.

Owen, M.J., Sawa, A., Mortensen, P.B., 2016. Schizophrenia. Lancet 388(10039), 86–97.

Palmer, J., Kreutz-Delgado, K., Makeig, S., 2016. AMICA: An Adaptive Mixture of Independent Component Analyzers with Shared Components. http://citeseerx.ist.psu.edu/viewdoc/summary?doi=10.1.1.295.1351.

Palmer, J., M.S., Kreutz-Delgado, K., Rao, B., 2008. Newton Method for the ICA Mixture Model, Proceedings of the 33rd IEEE International Conference on Acoustics and Signal Processing (ICASSP 2008) 1805–1808. https://sccn.ucsd.edu/~jason/icassp08.pdf

Palmer, S.M., Rosa, M.G., 2006. Quantitative analysis of the corticocortical projections to the middle temporal area in the marmoset monkey: evolutionary and functional implications. Cereb Cortex 16(9), 1361–1375.

Perez, V.B., Tarasenko, M., Miyakoshi, M., Pianka, S.T., Makeig, S.D., Braff, D.L., Swerdlow, N.R., Light, G.A., 2017. Mismatch Negativity is a Sensitive and Predictive Biomarker of Perceptual Learning During Auditory Cognitive Training in Schizophrenia. Neuropsychopharmacology 42(11), 2206–2213.

Piazza, C., Miyakoshi, M., Akalin-Acar, Z., Cantiani, C., Reni, G., Bianchi, A.M., 2016. An Automated Function for Identifying EEG Independent Components Representing Bilateral Source Activity, XIV Mediterranean Conference on Medical and Biological Engineering and Computing 2016. Springer International Publishing 105–109.

Pion-Tonachini, L., Kreutz-Delgado, K., Makeig, S., 2019. ICLabel: An automated electroencephalographic independent component classifier, dataset, and website. Neuroimage 198, 181–197.

Plechawska-Wojcik, M., Kaczorowska, M., Zapala, D., 2019. The artifact subspace reconstruction (ASR) for EEG signal correction. A comparative study, Information systems architecture and technology: proceedings of 39th international conference on information systems architecture and technology – ISAT 2018: part II. Springer International Publishing, 125–135.

Rapaport, M.H., Bazzetta, J., McAdams, L.A., Patterson, T., Jeste, D.V., 1996. Validation of the Scale of Functioning in Older Outpatients With Schizophrenia. Am J Geriatr Psychiatry 4(3), 218–228.

Rasser, P.E., Schall, U., Todd, J., Michie, P.T., Ward, P.B., Johnston, P., Helmbold, K., Case, V., Soyland, A., Tooney, P.A., Thompson, P.M., 2011. Gray matter deficits, mismatch negativity, and outcomes in schizophrenia. Schizophr Bull 37(1), 131–140.

Rissling, A.J., Miyakoshi, M., Sugar, C.A., Braff, D.L., Makeig, S., Light, G.A., 2014. Cortical substrates and functional correlates of auditory deviance processing deficits in schizophrenia. Neuroimage Clin 6, 424–437.

Salisbury, D.F., Polizzotto, N.R., Nestor, P.G., Haigh, S.M., Koehler, J., McCarley, R.W., 2017. Pitch and Duration Mismatch Negativity and Premorbid Intellect in the First Hospitalized Schizophrenia Spectrum. Schizophr Bull 43(2), 407–416.

Spencer, K.M., Salisbury, D.F., Shenton, M.E., McCarley, R.W., 2008. Gamma-band auditory steady-state responses are impaired in first episode psychosis. Biol Psychiatry 64(5), 369–375.

Stone, W.S., Giuliano, A.J., Tsuang, M.T., Braff, D.L., Cadenhead, K.S., Calkins, M.E., Dobie, D.J., Faraone, S.V., Freedman, R., Green, M.F., Greenwood, T.A., Gur, R.E., Gur, R.C., Light, G.A., Mintz, J., Nuechterlein, K.H., Olincy, A., Radant, A.D., Roe, A.H., Schork, N.J., Siever, L.J., Silverman, J.M., Swerdlow, N.R., Thomas, A.R., Tsuang, D.W., Turetsky, B.I., Seidman, L.J., 2011. Group and site differences on the California Verbal Learning Test in persons with schizophrenia and their first-degree relatives: findings from the Consortium on the Genetics of Schizophrenia (COGS). Schizophr Res 128(1-3), 102–110.

Swerdlow, N.R., Bhakta, S., Chou, H.H., Talledo, J.A., Balvaneda, B., Light, G.A., 2016. Memantine Effects On Sensorimotor Gating and Mismatch Negativity in Patients with Chronic Psychosis. Neuropsychopharmacology 41(2), 419–430.

Tada, M., Kirihara, K., Koshiyama, D., Fujioka, M., Usui, K., Uka, T., Komatsu, M., Kunii, N., Araki, T., Kasai, K., in press. Gamma-Band Auditory Steady-State Response as a Neurophysiological Marker for Excitation and Inhibition Balance: A Review for Understanding Schizophrenia and Other Neuropsychiatric Disorders. Clin EEG Neurosci.

Tada, M., Kirihara, K., Mizutani, S., Uka, T., Kunii, N., Koshiyama, D., Fujioka, M., Usui, K., Nagai, T., Araki, T., Kasai, K., 2019. Mismatch negativity (MMN) as a tool for translational investigations into early psychosis: A review. Int J Psychophysiol 145, 5–14.

Tada, M., Nagai, T., Kirihara, K., Koike, S., Suga, M., Araki, T., Kobayashi, T., Kasai, K., 2016. Differential Alterations of Auditory Gamma Oscillatory Responses Between Pre-Onset High-Risk Individuals and First-Episode Schizophrenia. Cereb Cortex 26(3), 1027–1035.

Takahashi, H., Rissling, A.J., Pascual-Marqui, R., Kirihara, K., Pela, M., Sprock, J., Braff, D.L., Light, G.A., 2013. Neural substrates of normal and impaired preattentive sensory discrimination in large cohorts of nonpsychiatric subjects and schizophrenia patients as indexed by MMN and P3a change detection responses. Neuroimage 66, 594–603.

Teale, P., Collins, D., Maharajh, K., Rojas, D.C., Kronberg, E., Reite, M., 2008. Cortical source estimates of gamma band amplitude and phase are different in schizophrenia. Neuroimage 42(4), 1481–1489.

Thomas, M.L., Green, M.F., Hellemann, G., Sugar, C.A., Tarasenko, M., Calkins, M.E., Greenwood, T.A., Gur, R.E., Gur, R.C., Lazzeroni, L.C., Nuechterlein, K.H., Radant, A.D., Seidman, L.J., Shiluk, A.L., Siever, L.J., Silverman, J.M., Sprock, J., Stone, W.S., Swerdlow, N.R., Tsuang, D.W., Tsuang, M.T., Turetsky, B.I., Braff, D.L., Light, G.A., 2017. Modeling Deficits From Early Auditory Information Processing to Psychosocial Functioning in Schizophrenia. JAMA Psychiatry 74(1), 37–46.

Thuné, H., Recasens, M., Uhlhaas, P.J., 2016. The 40-Hz Auditory Steady-State Response in Patients With Schizophrenia: A Meta-analysis. JAMA Psychiatry 73(11), 1145–1153.

Toyomaki, A., Kusumi, I., Matsuyama, T., Kako, Y., Ito, K., Koyama, T., 2008. Tone duration mismatch negativity deficits predict impairment of executive function in schizophrenia. Prog Neuropsychopharmacol Biol Psychiatry 32(1), 95–99.

Tsuchimoto, R., Kanba, S., Hirano, S., Oribe, N., Ueno, T., Hirano, Y., Nakamura, I., Oda, Y., Miura, T., Onitsuka, T., 2011. Reduced high and low frequency gamma synchronization in patients with chronic schizophrenia. Schizophr Res 133(1-3), 99–105.

Uhlhaas, P.J., Singer, W., 2010. Abnormal neural oscillations and synchrony in schizophrenia. Nat Rev Neurosci 11(2), 100–113.

Umbricht, D., Krljes, S., 2005. Mismatch negativity in schizophrenia: a meta-analysis. Schizophr Res 76(1), 1–23.

van Os, J., Kapur, S., 2009. Schizophrenia. Lancet 374(9690), 635–645.

Vierling-Claassen, D., Siekmeier, P., Stufflebeam, S., Kopell, N., 2008. Modeling GABA alterations in schizophrenia: a link between impaired inhibition and altered gamma and beta range auditory entrainment. J Neurophysiol 99(5), 2656–2671.

Wilson, T.W., Hernandez, O.O., Asherin, R.M., Teale, P.D., Reite, M.L., Rojas, D.C., 2008. Cortical gamma generators suggest abnormal auditory circuitry in early-onset psychosis. Cereb Cortex 18(2), 371–378.

Wynn, J.K., Sugar, C., Horan, W.P., Kern, R., Green, M.F., 2010. Mismatch negativity, social cognition, and functioning in schizophrenia patients. Biol Psychiatry 67(10), 940–947.

Yamasue, H., Yamada, H., Yumoto, M., Kamio, S., Kudo, N., Uetsuki, M., Abe, O., Fukuda, R., Aoki, S., Ohtomo, K., Iwanami, A., Kato, N., Kasai, K., 2004. Abnormal association between reduced magnetic mismatch field to speech sounds and smaller left planum temporale volume in schizophrenia. Neuroimage 22(2), 720–727.

